# In vivo measurement of the Young’s modulus of the cell wall of single root hairs

**DOI:** 10.1101/2022.12.20.521227

**Authors:** David Pereira, Thomas Alline, Atef Asnacios

**Author notes:** equal contribution (David Pereira, Thomas Alline).

## Abstract

Root hairs are cells from the root epidermis that grow as long tubular bulges perpendicular to the root. They can grow in a variety of mechanical or chemical environments. Their mechanical properties are mainly due to their stiff cell wall which also constitutes a physical barrier between the cell and its environment. Thus, it is essential to be able to quantify the cell wall mechanical properties and their adaptation to environmental cues. Here, we present a technique we developed to measure the Young’s (elastic) modulus of the root hair cell wall. In essence, using custom-made glass microplates as cantilevers of calibrated stiffness, we are able to measure the force necessary to bend a single living root hair. From these experiments one can determine the stiffness and Young’s modulus of the root hair cell wall.

## Introduction

Root hairs are differentiated cells that emerge from the root epidermis. They are highly elongated tubular structures of 10 μm in diameter and up-to few millimeters in length(1). These cells play a significant role in nutrients uptake and root-soil interactions. Indeed, they increase the surface of exchange of the root by up-to 2-fold (2). During their growth, RH cells invade soils displaying a huge diversity of mechanical properties (stiff, soft, dry, wet, heterogeneous, rocky, dense, etc.).

The main component ensuring RH structural integrity is the cell wall (CW). Due to its rigid structure, it maintains the cell integrity upon turgor pressure and also protects cells from mechanical stresses. Spatio-temporal regulation of its mechanical properties is also essential for cellular growth. Indeed, upon isotropic turgor pressure, the regulated process of secretion, synthesis and modification of the cell wall at the tip leads to polar (e.g. oriented) growth of the root hair(3). The CW is composed of two layers (4), called the primary (PW) and the secondary (SW) cell wall. They have different localizations, structures, compositions and roles. The PW is mostly made of cellulose, hemicellulose and pectin, with a random orientation of the cellulose microfibrils. The PW is present on the shank and tip of the cell (4) and, is essential for the tip growth (5). In contrast, the SW is localized only on the shank. It displays a different cellulose network with longitudinally oriented and densely packed microfibrils(4). The SW has a significant role in the strengthening of the RH.

Thus, measuring the mechanical properties of the RH CW, in particular at different stages of growth, or after growth in different environments, would be important to understand the complex process of cell growth, cell strengthening, and could also help to model the cell growth. Measurements of the Young’s modulus of the cell wall have been reported on many other tip growing systems (such as pollen tube, Hyphae, fission yeast) submitted to bending or buckling experiments (6)(7)(8). On root hair cells, only local measurements of the Young’s modulus and the stiffness have been carried out using AFM (9)(10).

Here we present a novel approach to quantify the mechanical properties of the root hair cell wall, using a microplate-based technique to measure the Young’s modulus of the cell wall through a bending experiment.

## Method and Results

### Measurement of the Young’s modulus of the Root hair cell wall

The root hair cell wall can be viewed as an elastic beam. For a bending experiment where a force F is applied at the tip of the RH perpendicularly to the RH axis, the Young’s modulus is given by:

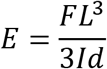

Where L is the RH length (more precisely the distance between RH base and the point of force application), d its deflection, and I the second moment of area of the beam.

In such an experiment, to have access to the Young’s modulus of the root hair cell wall, one has to know or measure all these parameters. With our experimental setup we can measure the force, the deflection, the length of the root hair and, thanks to previous studies, we can estimate the second moment of area.

Our experimental setup is a combination of a microfluidic-like system (MLS)(11) where *Arabidopsis thaliana* seedlings are grown in solid media, and a micromanipulation setup controlling a glass microplate used as a force sensor (cantilever of calibrated stiffness). On the one hand the MLS allows us to grow seedlings for weeks and to image them under a microscope. It also allows one to keep the main root close to the coverslip, and to minimize root movements during imaging and mechanical measurements, reducing thus noise and uncertainties.

On the other hand, the glass microplate is used to bend the RH cell. It was designed in size and shape to fit the range of forces required for the experiment. To do so, a microplate with a rectangular cross section was heated and pulled with a micropipette puller as previously described (12). The cantilever can be viewed as a spring with a characterized stiffness k. The parameters were set-up to have a microplate bending stiffness of about 50nN/μm.

For the measurements, a small quantity of the solid medium around the RH cells was removed and replaced by liquid media. The whole system was then placed under the microscope, the calibrated glass microplate was immersed and, thanks to micromanipulators, placed close to the root hair tip.

Then, piezoelectric actuators are used to move the microplate base in order to deflect the RH. Thus, knowing the imposed displacement D of the microplate base and the RH deflection d (measured on the image), the force F applied on the root hair is given by: F = k(D – d). The length L of the RH is measured by taking the distance between the position where the force is applied and the RH base (figure 1B). Because F, L and d are measured parameters, one can compute, without any assumption, the value of the product of the Young’s modulus by the second moment of area EI which is a good characteristic of the mechanical structure of the RH (figure 1D-E). For 20 Arabidopsis RH of an average length of 189 +/− 19 μm, we found an average value of EI of 6.1e-14 +/− 1.1e-14 N.m^2^.

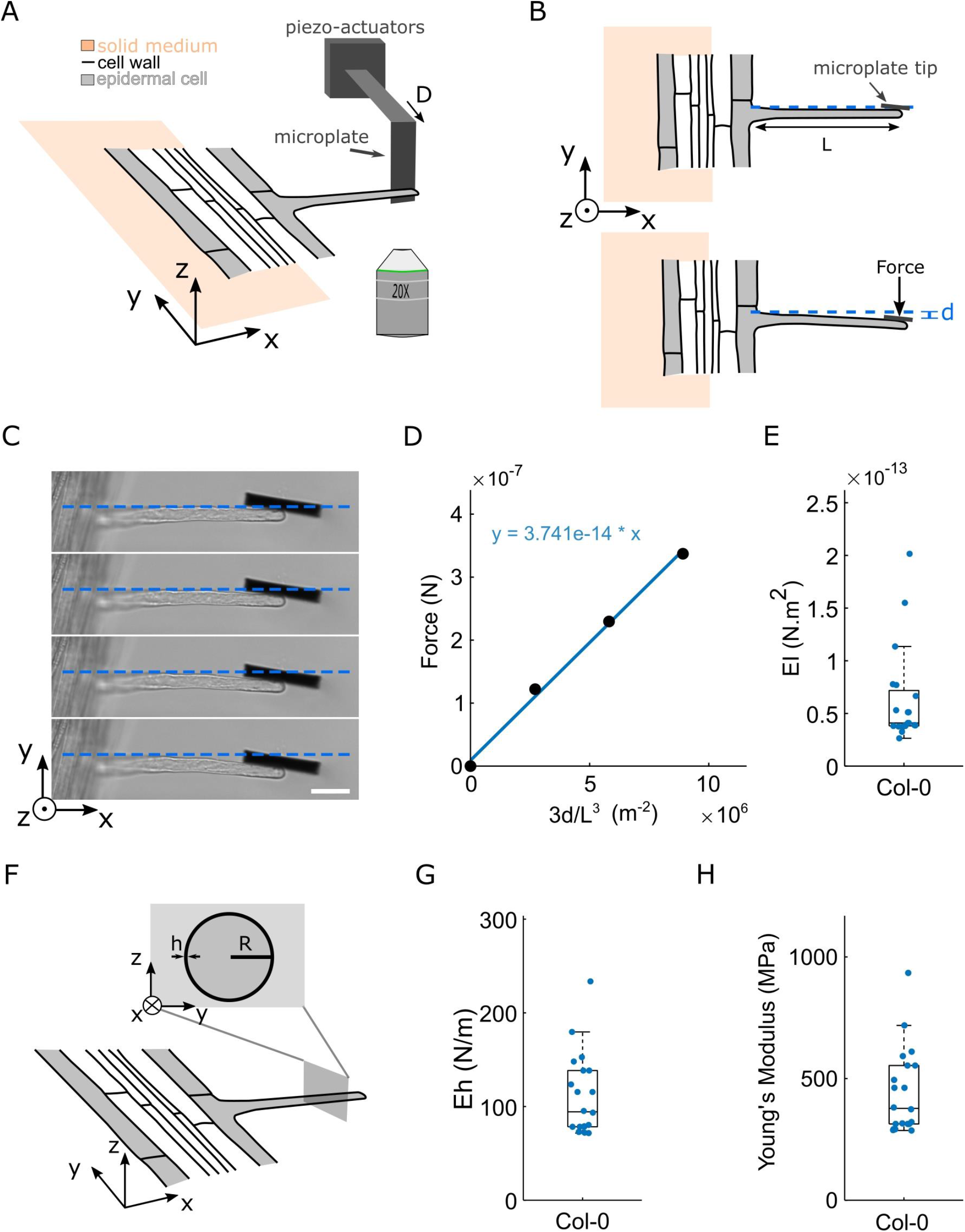
(A) Schematic representation of the experiment. The microplate is placed vertically against the tip of the RH. Then, thanks to a piezo-actuator, the microplate base is moved by a distance D resulting in the bending of the root hair. (B) Schematic representation in the XY plane. Top, before the measurement. Bottom, the root hair is deflected by a length d. The microplate deflection is thus (D-d) and it exerts a force F= k (D-d) on the root hair, with k the stiffness of the microplate. (C) Sequence of images displaying deflection increments applied on a single RH. Between each image the base of the microplate is moved by 5 μm. (scale bar=30μm). (D) data corresponding to (C). The force exerted is expressed as a function of 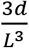. The blue curve represents the linear fit of the data. Its slope y represents the factor EI. (E) Boxplot showing the distribution of the measured factor EI for n=20 root hairs. (F) Schematic representation of a root hair viewed as a hollow cylinder of radius R and negligible width h. (H) Boxplot showing the distribution of the stiffness Eh (n=20) (I) Boxplot showing the distribution of the cell wall Young Modulus E when h is taken as equal to 250μm (n=20).

The RH cell has a tubular structure with a thin cell wall thickness which is negligible as compared to the RH diameter. Thus, its second moment of area can be expressed as:

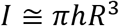

With h the cell wall thickness and R the radius of the root hair.

By measuring the root hair radius R one can thus have access to an effective stiffness Eh characteristic of the CW, and independent of R and L, e.g. independent of the shape of the RH. For the 20 RH tested, we found an average CW effective stiffness of 110 +/− 9 N/m (figure 1G).

Using electron microscopy, Akkerman et al(4) measured the cell wall thickness of root hair cells, and found a value around 250nm. Assuming h=250 nm, one can then estimate the cell wall Young’s modulus E. We found E = 443 +/− 39 MPa.

In sum, we developed a novel technique allowing one to measure the effective stiffness and the Young’s modulus of the cell wall of living root hair cells.

## Discussion

The cell wall is an essential component of root hair cells. Measuring its mechanical properties would provide a better understanding of the cell growth and its possible adaptation to different chemical or mechanical conditions.

The Young’s modulus of biological systems such as pollen tube, Hyphae and fission yeast have been previously measured (6)(7)(8). Due to the complexity in shape, size and of the culture condition, different techniques were developed to allow one to measure the mechanical properties of each of these biological systems. Here we describe a method to measure the Young’s modulus of root hair cells using a microplate-based technique. Our method is well adapted for measurements on single living RH of different sizes, or at different growth stages, and it also gets rid of the movement of the root.

The reported value of the Young’s modulus of pollen tube, using fluid flow to bend the cell, was 350 MPa (6). Interestingly, this value is relatively close to the one we found for the RH cells. In contrast, measurements of RH cell wall modulus using AFM showed values much lower (1-7MPa) compared to the one presented in this study (9)(10). This could be explained by the technique itself which permits only local measurements of the mechanical properties(13).

During growth, root hair cells invade a diversity of chemical and mechanical environments that can affect the cell growth(14) (15). Thus, it would be interesting to measure the mechanical properties of RH cells that were grown in this variety of environments.

Such mechanical measurements could also help understanding the biochemical pathways controlling CW composition and properties. For instance, it has been reported that the cell wall composition is controlled by the ERULUS kinase. ERULUS mutants are shorter, larger and present defects in their cell wall. Thus, it would also be interesting to measure the mechanical properties of the ERULUS mutant cell wall.

Along this line, one important point to notice is that root hair cells have a primary and a secondary cell wall(16) but the interplay between them is still unclear. Here we measure the young modulus of the two cell walls combined. The secondary cell wall rigidifies the root hair cell shank but it is still unclear when this phenomenon starts. Therefore, looking at the mechanical properties of the cell wall at different time points, during growth, could help to clarify the temporality of the secondary cell wall synthesis.

Moreover, we could also measure the mechanical properties of the RH by applying the force not only at the tip but at different positions along the RH long axis to quantify cell wall heterogeneities. Indeed, it has recently been shown that there could be some localized cell wall remodelling in the shank of root hairs(17).

Of note, while we have here used an estimation of the cell wall thickness taken from previous studies, future work, in particular on specific mutants, will imply to determine the CW thickness in parallel to the mechanical measurements, for instance through electron microscopy or fluorescence techniques.

In conclusion, we developed a novel microplate-based technique to measure the mechanical properties of the cell wall of RH cells, allowing to enhance our understanding of the cell wall mechanics and, consequently, the cell growth.

## Acknowledgements

We thank Herman Höfte and Sebastjen Schonaers for fruitful discussions and for providing the the col-0 lines. This Work was supported by the Centre National De la Recherche Scientifique (CNRS) and by HFSP grant 2018, RGP, 009. The study was partially supported by the labex “Who AM I?”, labex ANR-11-LABX-0071, as well as the Université Paris Cité, Idex ANR-18-IDEX-0001, funded by the French Government through its “Investments for the Future” program and also by “Mecha-Nuc” project ANR-20CE13-0025-03.

## Competing interests

The authors declare no competing interests.

## Author contributions

DP and AA developed the conceptual framework and designed the study. AA sought funding.

DP and TA performed the experiments and the quantitative analyse of the results.

DP, TA and AA analysed the findings and wrote the manuscript.

## Notes

### Competing Interest Statement

The authors have declared no competing interest.

